# High-Density Wild-Type IL-2 Nanoparticles Preferentially Enhance CD8⁺ T-Cell Expansion and Reprogram the Tumor Microenvironment

**DOI:** 10.64898/2026.07.14.738558

**Authors:** Ruijie Wang, Pramod Kumar, Noah A. Crumrine, Tongchatra Watcharawittayakul, Alyssa Wallstrum, Moataz Reda, Gordon Mills, Worapol Ngamcherdtrakul, Wassana Yantasee

## Abstract

Low response rates to immune checkpoint inhibitors (ICIs) in solid tumors are often driven by insufficient tumor-infiltrating CD8⁺ T cells and immunosuppressive tumor microenvironment (TME). Although interleukin-2 (IL-2) potently expands and activates CD8⁺ T cells, its clinical use is limited by rapid clearance, dose-limiting toxicity, and regulatory T cell (T_reg_) stimulation. Engineered IL-2 variants have not yet achieved meaningful clinical efficacy. Here, polymer-modified mesoporous silica nanoparticles displaying dense, unmodified wild-type IL-2 on their surface (IL2-NP) are developed, conferring proteolytic stability and tumor retention. IL2-NP enables avidity-mediated CD8⁺ T cell binding and enhances proliferation and effector function without increased T_reg_ binding or proliferation.

Intratumoral IL2-NP expands CD8⁺ T cells, increases CD8⁺/T_reg_ ratios, and reprograms TME through dendritic cell activation and M1-like macrophage polarization. IL2-NP induces regression of both treated and untreated distant colorectal tumors in a CD8⁺ T cell-dependent manner. IL2-NP synergizes with ICIs and leads to complete tumor regression and immunological memory that protect against rechallenge. Treatment is well tolerated, with strong efficacy also observed in triple-negative breast and metastatic ovarian cancer models.

Overall, intratumoral IL2-NP elicits robust systemic antitumor immunity, offering a promising strategy to enhance ICIs, cancer vaccines, and adoptive T-cell therapies.

**Graphical abstract:** This work introduces a nanoparticle platform that overcomes major shortcomings of IL-2 immunotherapy by presenting wild-type IL-2 at high density on the nanoparticle surface, thereby increasing binding avidity to effector T cells. The resulting IL-2 nanoparticles enhance cytotoxic T cell expansion, reprogram the tumor microenvironment, and augment responses to immune checkpoint blockade to achieve robust ant-tumor immune response in mouse tumor models.

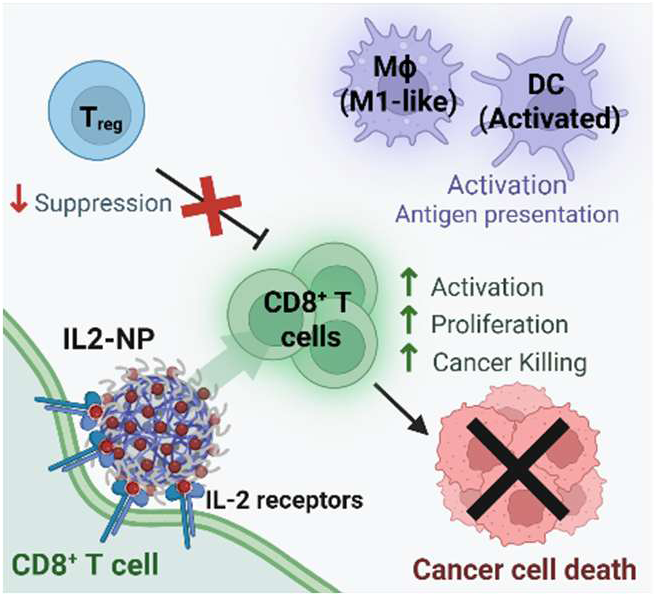

## 1. Introduction

Immune checkpoint inhibitors (ICIs) have transformed cancer therapy. However, while ICIs have generated durable responses in some patients,^1–3^ many tumor types remain poorly responsive, in part due to factors such as insufficient tumor-reactive T cell infiltration and immunosuppressive features in the tumor microenvironment.^4–10^ This has motivated continued interest in approaches that expand tumor-reactive T cells.^11^

Interleukin-2 (IL-2) is a potent T cell growth factor that has been explored for cancer immunotherapy. Aldesleukin, a recombinant human IL-2, is FDA-approved as monotherapy for the treatment of metastatic melanoma and renal cell carcinoma.^12^ IL-2 prefers to activate regulatory T cells (T_reg_), which express high-affinity trimeric IL-2 receptors (IL-2Rαβγ).^13^ In contrast, resting CD8^+^ T cells predominantly express intermediate-affinity dimeric IL-2Rβγ, which requires substantially higher IL-2 concentrations for activation.^13,14^ This differential receptor affinity provided the original rationale for high-dose regimens, where dosage sufficient to activate CD8^+^ T cells would be required due to their lower receptor affinity in the resting state.^14,15^ Even with such a regimen, the response rate to aldesleukin in metastatic melanoma and renal cell carcinoma is only 10-15%,^15^ and severe systemic toxicity, such as vascular leak syndrome, further limits its clinical use.^15^ Therefore, there is a need for a new IL-2 strategy that stimulates CD8^+^ T cells without accompanying dosing burden and systemic toxicity.

Several strategies have been explored to improve the therapeutic index of IL-2. Among these, non-α IL-2 variants, designed to avoid binding to the alpha subunit of the IL-2 receptor on T_reg_, have emerged as a major class of IL-2-based agents in clinical development.^14^ Some variants also incorporate PEGylation or Fc fusion to improve the pharmacokinetic profile. However, these approaches do not enhance engagement with IL-2Rβγ on CD8^+^ T cells, and several have shown disappointing clinical results, leading to the discontinuation of their development.^16–19^ These results suggest that avoiding IL-2Rα binding or simply extending circulation half-life may be insufficient to generate durable anti-tumor immunity. In contrast to previous understanding that CD8^+^ T cells do not express IL-2Rα, emerging evidence has shown that IL-2Rα is upregulated specifically in tumor-infiltrating CD8^+^ T cells^20^ and activated CD8^+^ T cells^21^, a finding consistent with a report showing that wild-type (WT) IL-2 outperforms non-α IL-2 variants in several mouse tumor models.^21^ Together, these findings suggest that preserving WT IL-2 biology, rather than eliminating IL-2Rα engagement, may be important for maximizing CD8^+^ T cell anti-tumor responses.

This study employs a polymer-coated mesoporous silica nanoparticle platform (Pdx-NP^TM^)^22^ that presents WT IL-2. We hypothesized that dense surface presentation of IL-2 would promote multivalent interactions with intermediate-affinity IL-2 receptors on CD8^+^ T cells, increasing binding avidity and enhancing CD8^+^ T-cell responses without increasing T_reg_ activation compared with free IL-2. By presenting wild-type IL-2, the platform would also preserve engagement of IL-2Rα on activated and tumor-infiltrating CD8^+^ T cells, a feature absent in non-α IL-2 variants. To test our hypothesis, we evaluated the effect of IL-2 surface density on T cell proliferation, examined the activities of the optimal IL-2 nanoparticle formulation in vitro, and assessed its therapeutic efficacy and treatment-associated immune changes in mouse tumor models.

## 2. Results

### 2.1. Synthesis and characterization of IL2-NP

We utilized our patented polymer-coated mesoporous silica nanoparticle platform (NP) to deliver wild-type IL-2. Briefly, 50-nm mesoporous silica nanoparticles were sequentially coated with crosslinked PEI and PEG similar to our prior work,^23^ and recombinant mouse IL-2 was then loaded through electrostatic interaction with PEI (**Figure 1A**), thereby avoiding potential conjugation-driven conformational change. An outer PEG layer prevents nanoparticle aggregation and protects IL-2 from enzymatic degradation without interfering with IL-2 receptor binding. The final polymer composition was 23.4% PEI and 20.9% PEG by weight of NP, as assessed by Thermogravimetric Analysis (TGA).

**Figure 1.**
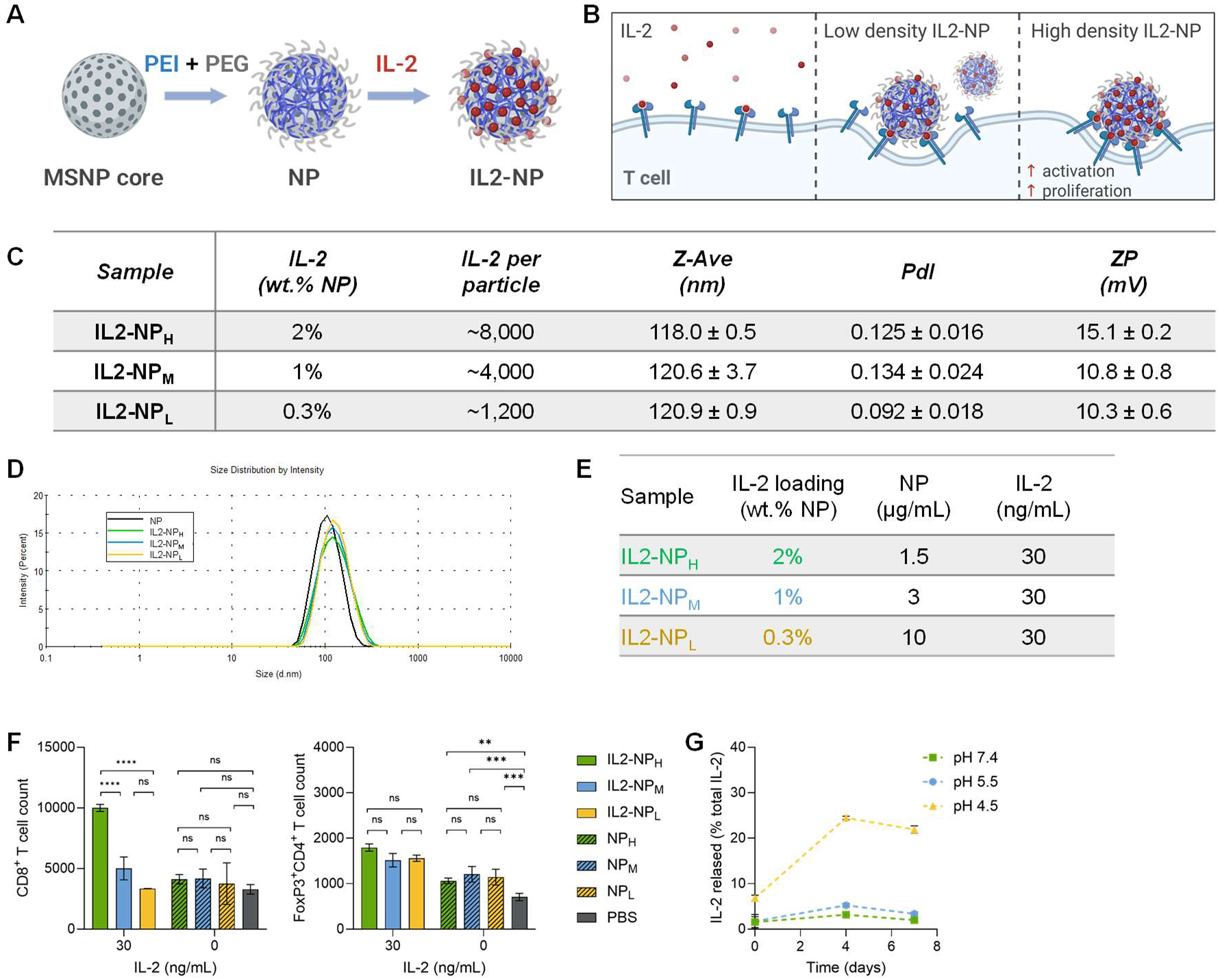
Effect of IL-2 surface density on T cell proliferation. (**A**) Step-by-step synthesis of the polymer coated MSNP (NP) and IL-2 loaded NP (IL2-NP). (**B**) Proposed binding mechanism. Free IL-2 and low-density IL2-NP bind weakly to the intermediate affinity dimeric IL-2Rβγ receptors on CD8^+^ T cells, whereas high-density IL2-NP is hypothesized to enhance receptor binding avidity through multi-receptor engagement on CD8^+^ T cells. (**C**) Composition of IL-2 (by weight and by number of IL-2 molecules per particle), size, Polydispersity Index (Pdl), and zeta potential of NP loaded with high, medium, and low density of IL-2 (IL2-NP_H_, IL2-NP_M_, IL2-NP_L_, respectively). (**D**) Hydrodynamic size distribution of NP, IL2-NP_H_, IL2-NP_M_, and IL2-NP_L_. (**E-F**) Effect of IL-2 loading density on the proliferation of CD8^+^ T cells and T_reg_. (**E**) In vitro treatment concentrations. (**F**) Proliferation of CD8^+^ T cells (left) and T_reg_ (CD4^+^FoxP3^+^, right) from isolated mouse T cells (1 x 10^5^ cells) after 48-hour treatment with IL2-NP_H_, IL2-NP_M_, or IL2-NP_L_ at a matching dose of 30 ng/mL IL-2, as well as nanoparticle control (no IL-2) at corresponding doses as shown in (**E**). (**G**) Release profile of IL-2 from IL2-NP incubated in PBS (pH 7.4), or 100 mM sodium acetate buffers (pH 4.5 and pH 5.5) after 0-7 days of incubation at 37°C. Data presented as mean ± SD from 2 replicates. (Figure 1A-B Created with BioRender.com).

Higher IL-2 surface density is hypothesized to increase binding avidity on CD8^+^ T cells (**Figure 1B**). We first assessed whether increasing the loading density of IL-2 on the nanoparticle would alter the balance of activity between CD8^+^ T cells and T_reg_. Nanoparticle formulations with high, medium, and low density of IL-2 (IL2-NP_H_, IL2-NP_M_, IL2-NP_L_, respectively) were prepared by adding different amounts of IL-2 to the nanoparticle suspension in PBS. All three formulations achieved near-complete loading efficiency (>95%), as determined by BCA protein assay (**Figure 1C**). Further increases in IL-2 concentration did not result in higher loading beyond 2 wt.%. The resulting IL2-NPs showed hydrodynamic size of around 120 nm and a mild positive charge of 10-15 mV (**Figures 1C-D**).

We next assessed the bioactivity by treating isolated mouse T cells with each IL2-NP formulation at a matched final IL-2 concentration of 30 ng/mL, and quantifying expansion of different T cell subsets after 48 hours by flow cytometry (**Figures 1E-F**). Among the IL-2 loaded formulations, high-density IL2-NP_H_ (2 wt.%) induced significantly greater CD8^+^ T cell expansion compared to the IL2-NP_M_ and IL2-NP_L_ (**Figure 1F, left**). In contrast, all three formulations resulted in similar levels of T_reg_ expansion (**Figure 1F, right**). Dose-matched NP controls (without IL-2) showed no significant effect on cell count at the tested concentrations (**Figure 1F**). Based on these results, IL2-NP_H_, containing 2 wt.% IL-2 (approximately 8,000 IL-2 molecules per particle, **Figure 1C**), was selected for subsequent experiments and is referred to as IL2-NP hereafter.

To take advantage of binding avidity, IL-2 must remain associated with the nanoparticle during interaction with T cells. We therefore measured IL-2 release from the nanoparticles under different buffer conditions mimicking physiological condition (PBS, pH 7.4), mildly acidic condition (sodium acetate buffer pH 5.5) similar to the tumor microenvironment,^24^ and more acidic endosomal/lysosomal pH (sodium acetate buffer pH 4.5). Over the course of seven days at 37°C incubation, IL-2 remained associated with nanoparticles at pH 7.4 and pH 5.5, whereas greater release was observed under more acidic pH 4.5 (**Figure 1G**). The PEI polymer buffers in the pH 5.5–7.4 range, but becomes protonated at pH 4.5, leading to reduced electrostatic interactions with IL-2 and facilitating its release. Together, these data established a high-density IL2-NP with favorable physicochemical properties for subsequent in vitro and in vivo testing.

### 2.2. Functional activities of IL2-NP in vitro

We next tested whether the optimal IL2-NP formulation enhance IL-2 properties and activities including T cell proliferation, resistance to proteolytic degradation, and T cell engagement and uptake.

To quantify biological activity of IL2-NP versus free IL-2, we performed a proliferation assay using CTLL-2, an IL-2-dependent mouse T cell line.^25^ Proliferation of these cells upon IL2-NP vs. IL-2 treatments was measured by CTG assay. IL2-NP promoted CTLL-2 proliferation with an approximately 12-fold lower EC_50_ (half maximal effective concentration) compared to free IL-2 (7.88 ng/mL vs. 0.65 ng/mL, respectively; **Figure 2A**). In addition to CTLL-2 cell line, enhanced proliferation by IL2-NP over free IL-2 was also observed in primary mouse T cells (**Figure 2B**).

**Figure 2.**
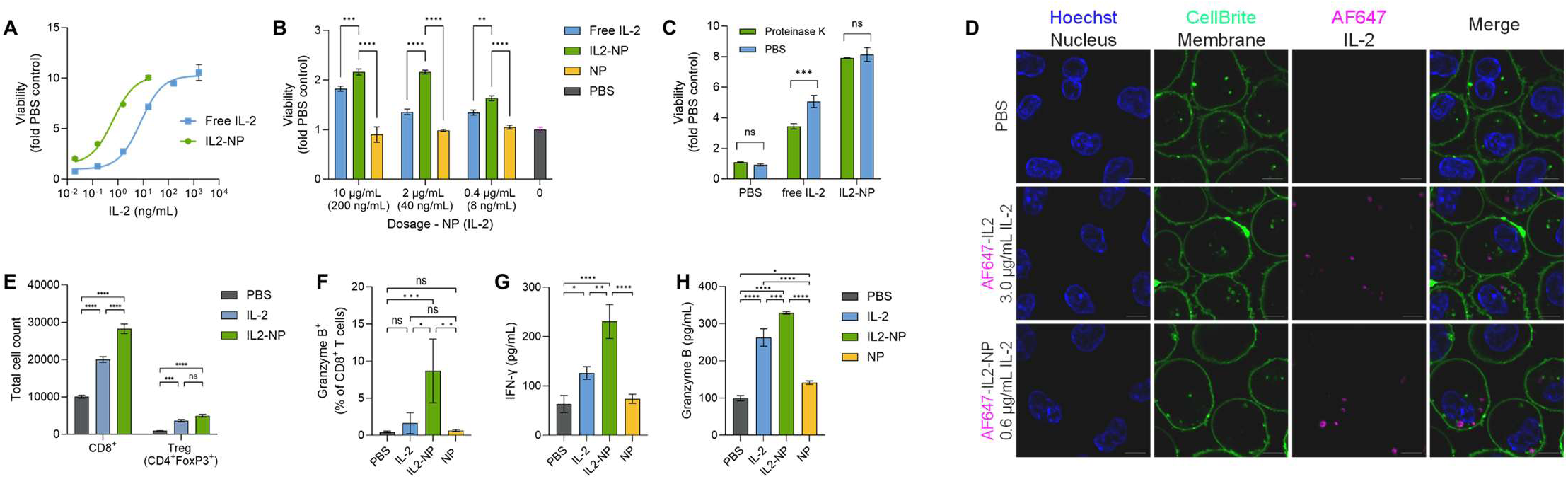
Mechanistic characterization of IL2-NP. (**A**) Proliferation of CTLL-2 cells as a function of IL-2 dose, delivered as free IL-2 or IL2-NP, measured after 48 hours using the CTG viability assay. (**B**) Viability of mouse splenic T cells treated with IL-2, NP, and IL2-NP for 48 hours. (**C**) Proliferation of murine splenic T cells treated with free IL-2 or IL2-NP (200 ng/mL IL-2), pre-incubated with 5 µg/mL proteinase K or PBS for 10 min at 37°C. (**D**) Confocal imaging of AF647-IL2-NP vs. AF647-IL2 after 1 hour incubation with CTLL-2 cells. Scale bars: 5 µm. (**E**) Flow cytometry analysis of CD8^+^ T cells and T_reg_ (CD4^+^FoxP3^+^) cell counts in mouse splenic T cells (5 x 10^5^ cells) treated with free IL-2 or IL2-NP (both with 30 ng/mL IL-2) for 24 hours. (**F**) Flow cytometry analysis of Granzyme B^+^ CD8^+^ T cells (CD3^+^CD8^+^) from mouse splenic T cells. (**G-H**) Secretion of IFN-γ (**G**) and Granzyme B (**H**) by ELISA in cell culture media of activated mouse splenic T cells co-cultured with LLC-JSP lung cancer cells (10:1 ratio), following 48-hour treatment with IL-2 or IL2-NP (200 ng/mL IL-2). Viability, flow cytometry and ELISA data presented as mean ± SD from 3 independent replicates.

Since proteases are commonly upregulated in the tumor microenvironment^26^ and can degrade cytokines like IL-2, we next tested whether the nanoparticles protect IL-2 from proteolytic degradation. Free IL-2 and IL2-NP were subjected to accelerated degradation with 5 µg/mL proteinase K or PBS for 10 min at 37°C, followed by evaluation of T-cell proliferative activity relative to untreated controls (no proteinase treatment). While the protease treatment significantly reduced the activity of free IL-2, it did not change IL2-NP activity (**Figure 2C**).

These results indicate that the nanoparticle formulation protects IL-2 from enzymatic degradation, likely through steric shielding by the PEG layer, consistent with our previous findings demonstrating siRNA protection using the same nanoparticle platform.^22^

Next, the interaction of IL2-NP with T cells was evaluated. Previous reports suggest that IL-2-receptor complex is internalized following T cell receptor engagement,^27^ which is required for downstream signaling.^28,29^ Alexa Fluor 647 (AF647)-labeled IL-2 was used to assess the interaction of IL-2 versus IL2-NP with CTLL-2 T cells by confocal microscopy. Both free AF647-IL2 and AF647-IL2-NP were internalized by T cells. However, AF647-IL2-NP produced stronger fluorescent signals in CTLL-2 cells compared to free AF647-IL2 under the tested conditions (**Figure 2D**). Notably, free AF647-IL2 required a 5-fold higher dose than AF647-IL2 on NP to achieve detectable signal in T cells by confocal microscopy.

To determine the effect of IL2-NP versus free IL-2 on proliferation of CD8^+^ T cells and T_reg_, mouse splenic T cells were treated with free IL-2 or IL2-NP, followed by flow cytometry analysis. IL2-NP treatment enhanced the expansion of CD8^+^ T cells while maintaining similar T_reg_ expansion, compared to free IL-2 (**Figure 2E**).

In addition to enhancing CD8^+^ T cell proliferation, IL2-NP also enhanced the cytotoxic activity of T cells. CD8⁺ T cell-mediated cancer cell killing relies on effector molecules such as IFN-γ and granzyme B.^30,31^ **Figure 2F** shows that IL2-NP increased the proportion of granzyme B^+^ among CD8^+^ T cells compared to free IL-2 or PBS and NP controls. To assess whether IL2-NP enhanced T cell effector functions in a T cell–cancer cell co-culture system, activated mouse T cells were co-cultured with LLC-JSP cancer cells and treated with free IL-2 or IL2-NP. IL2-NP treatment resulted in higher levels of IFN-γ (**Figure 2G**) and granzyme B (**Figure 2H**) production in the cell culture media compared to free IL-2, while NP alone had little effect compared to PBS.

Together, these findings show that IL2-NP imparts many desirable properties. IL2-NP enhanced CD8⁺ T cell proliferation without increasing T_reg_ proliferation compared with free IL-2, preserved IL-2 activity after protease exposure, increased T cell uptake, and enhanced CD8^+^ T cell effector function as reflected by granzyme B and IFN-γ production.

### 2.3. Local IL2-NP treatment induces systemic anti-tumor response and immune memory in a bilateral colorectal cancer model

We hypothesized that intratumoral IL2-NP delivery could generate systemic antitumor immunity. A bilateral MC38 colorectal tumor model was used to assess whether the intratumoral IL2-NP treatment of one tumor could affect the growth of an untreated contralateral (distant) tumor, demonstrating systemic antitumor activity. The model was also used to assess the dependence of CD8^+^ T cells and immune memory response associated with the treatment.

First, we evaluated whether locally administered IL2-NP remained within the injected tumor. We administered free IL-2 versus IL2-NP (using AF680-label IL-2) intratumorally and harvested tumors to assess tumor retention by monitoring fluorescence intensity of AF680 (**Figure 3A**). No detectable signal was observed in the distal tumor one day after injection (**Figure 3B**), indicating minimal leakage of IL-2 from the treated tumor. Consistent with this observation, in a separate distribution experiment using NP loaded with dye-labeled siRNA, signal remained localized in tumor for up to 10 days, which increased accumulation (AUC) by 8-fold compared to free siRNA (**Figure 3C**). These data support enhanced tumor retention of small molecules such as IL-2 and siRNA through nanoparticle delivery following intratumoral administration. Notably, siRNA and IL-2 have similar molecular weight (14 kDa for siRNA and 17 kDa for IL-2). Both are loaded onto the nanoparticle platform via electrostatic interactions and share similar delivery requirements, including protection against enzymatic degradation and rapid renal clearance.

**Figure 3.**
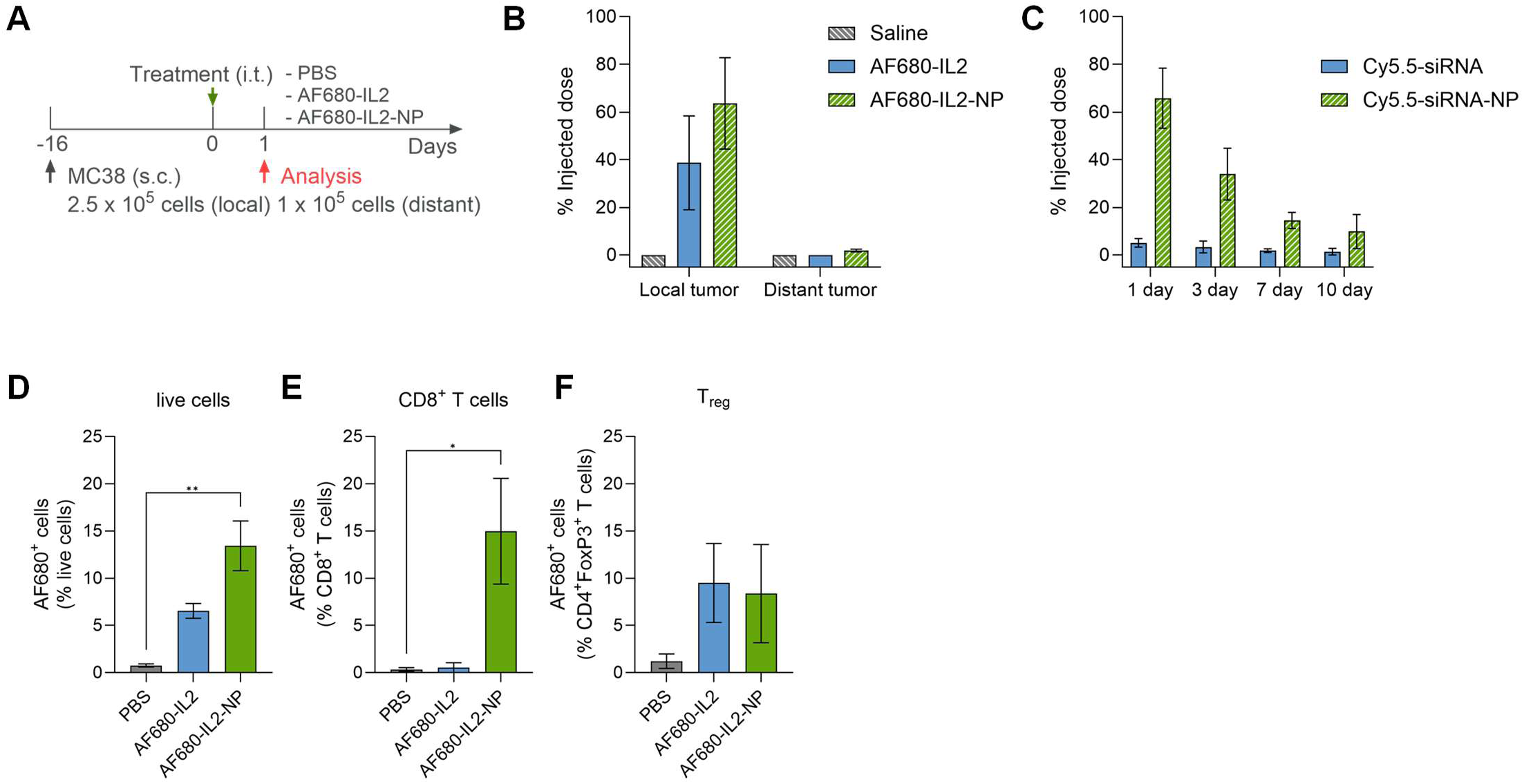
Enhanced tumor retention and preferential CD8^+^ T-cell binding of IL2-NP. (**A-B**) Tumor retention of IL-2 delivered by the nanoparticle platform. (**A**) Study timeline. Mice bearing bilateral MC38 tumors (n=3-4) were treated with intratumoral injections of saline, free IL-2 or IL2-NP (using AF680-labeled IL-2) and analyzed one day later. (**B**) Retention of IL-2 in the treated and distant tumors reported as percentage of injected dose. (**C**) Mice bearing B16F10 tumors were treated i.t. with free siRNA or siRNA-NP (using Cy5.5-tagged siRNA). Tumors were harvested on day 1, 3, 7, and 10 days post injection (n=2-4). The percentage of injected dose was analyzed similar to (**B**). (**D-F**) Tumors from (**A**) were harvested 24 hours after free IL-2 or IL2-NP treatment, followed by flow analysis of IL-2 positive (via AF680^+^) cells and reported as percentage of live cells (**D**), CD8^+^ T cells (**E**), and T_reg_ (**F**). Data (n=3-4) presented as mean ± SEM.

To understand the binding of IL2-NP versus free IL-2 on various immune cell populations in the tumor, we also analyzed the tumors at 24 hours post treatments with flow cytometry (**Figure 3A**). AF680-IL2-NP treatment resulted in a significantly higher proportion of AF680-IL2^+^ live cells in the tumor compared to free AF680-IL2 treatment (**Figure 3D**). Further analysis of T-cell subsets revealed that AF680-IL2-NP preferentially enhanced association with CD8^+^ T cells, compared to free AF680-IL2, which showed levels comparable to saline control (**Figure 3E**). In contrast, AF680-IL2-NP did not increase the association with T_reg_, compared to free AF680-IL2 (**Figure 3F**).

Once we confirmed that IL2-NP retained IL-2 in locally injected tumor, the antitumor efficacy of IL2-NP in both treated and distal tumors was assessed following the treatments as shown in **Figure 4A**. The MC38 cell line used in this study was obtained from Kerafast (originating from an NCI/NIH clone) and was reported to be less immunogenic than the more commonly used Leiden University clone and lacks several neoantigens (i.e. mutated Rpl18 and Adpgk).^32^ IL2-NP significantly improved tumor control of both local and distal tumors compared to free IL-2. It achieved complete regression in a subset of mice (**Figures 4B–C**), whereas free IL-2 did not result in any cure. In this model, immune checkpoint blockade against PD-1 and CTLA-4 (ICIs) produced complete response in 2 of 7 mice, whereas IL2-NP combined with ICIs achieved 100% cure rate (**Figure 4C**).

**Figure 4.**
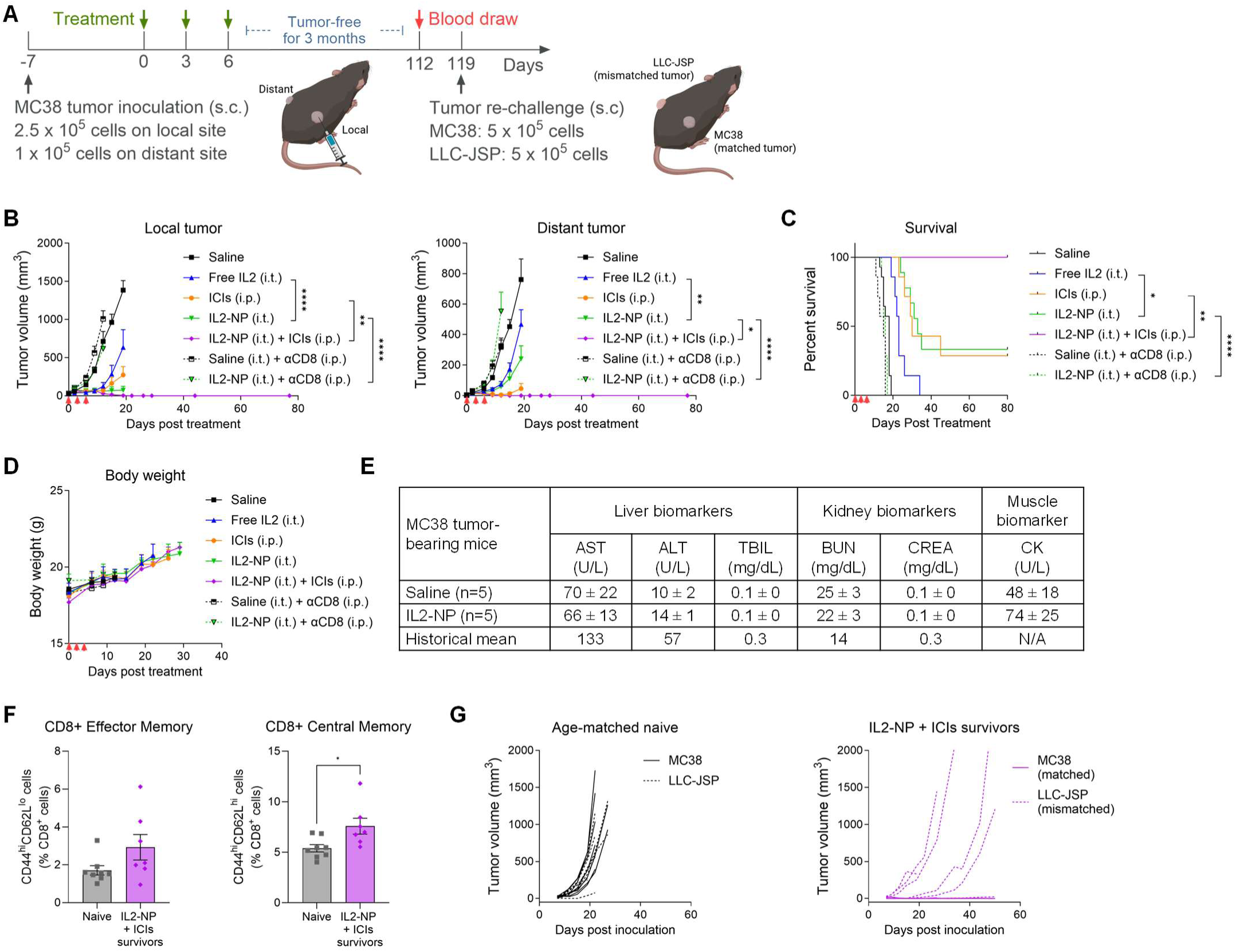
Efficacy and CD8^+^ T-dependence of IL2-NP treatment in a mouse model of MC38 colorectal cancer. (**A**) Schematic illustration of efficacy and immune memory study timeline. n = 7-14 per group from two independent experiments. (**B**) Growth curves of the local treated and the distant tumors. (**C**) Kaplan–Meier Survival curve. (**D**) Body weight of the mice. (**E**) Serum chemistry from bilateral MC38 tumor-bearing mice treated with three i.t. doses of IL2-NP or saline. Serum samples were collected at 1 day post last dose; data are compared with historical data from 8-10 week tumor-free female C57BL/6NCrl (n=141 to 161).^33^ (**F**) Circulating effector memory (CD44^hi^CD62L^lo^) (left) and central memory (CD44^hi^CD62L^hi^) T cells (right) in mice treated with IL2-NP + ICIs that remained tumor-free for 3 months, compared to those in age-matched naïve controls (n = 7-8). (**G**) MC38 (solid line) and LLC-JSP (dashed line) tumor growth in age-matched naïve mice and IL2-NP+ICIs treated survivors.

Next, we tested whether the antitumor efficacy of IL2-NP depended on CD8^+^ T cells. Mice were treated with a CD8-depleting antibody, followed by intratumoral IL2-NP or saline as described in **Figure 4A**. CD8 depletion did not significantly alter the tumor growth and survival compared to saline-treated controls. In contrast, the CD8 depletion completely abrogated the therapeutic effect of IL2-NP (**Figures 4B–C**), indicating that the antitumor activity is dependent on CD8^+^ T cells, as expected.

We also evaluated the tolerability of the dosing regimen used in this study. Repeated IL2-NP administration was associated with stable body weight (**Figure 4D**) and serum biochemistry values that did not indicate treatment-associated abnormalities in liver, kidney and muscle function (**Figure 4E**).

To determine whether the complete responses from IL2-NP and ICIs combination treatment were associated with durable immune memory, the treated mice that remained tumor-free for three months were evaluated for memory T cells in their peripheral blood and then rechallenged with matched and mismatched cancer cells. Elevated numbers of CD8^+^ effector memory and central memory cells (**Figure 4F**) were observed in the IL2-NP + ICIs treated mice compared to age-matched naïve controls. Upon rechallenge, naïve mice developed both matched MC38 and mismatched LLC-JSP tumors, whereas previously cured mice rejected MC38 tumor but not LLC-JSP tumors (**Figure 4G**). These results indicate that the combination treatment induced tumor-specific immune memory in mice that protected against tumor rechallenge. Mice cured by ICIs alone (from **Figure 4C**) did not exhibit any protection against tumor rechallenge (albeit with a small sample size).

### 2.4. IL2-NP treatment preferentially targets CD8^+^ T cells and reshapes the tumor microenvironment

To understand the effect of IL2-NP on various immune cell populations that contributed to the anti-tumor efficacy, we assessed immune changes by profiling lymphoid and myeloid compartments in tumors and tumor-draining lymph nodes by flow cytometry (**Figure 5A**). IL2-NP treatment significantly increased the proportion of CD8^+^ T cells in the tumors (**Figure 5B**), without increasing T_reg_ (**Supplementary Figure S1A**). Consequently, IL2-NP treatment resulted in a significant increase in CD8/T_reg_ ratio in both local and distant tumors compared to free IL-2 and saline controls (**Figure 5C**).

**Figure 5.**
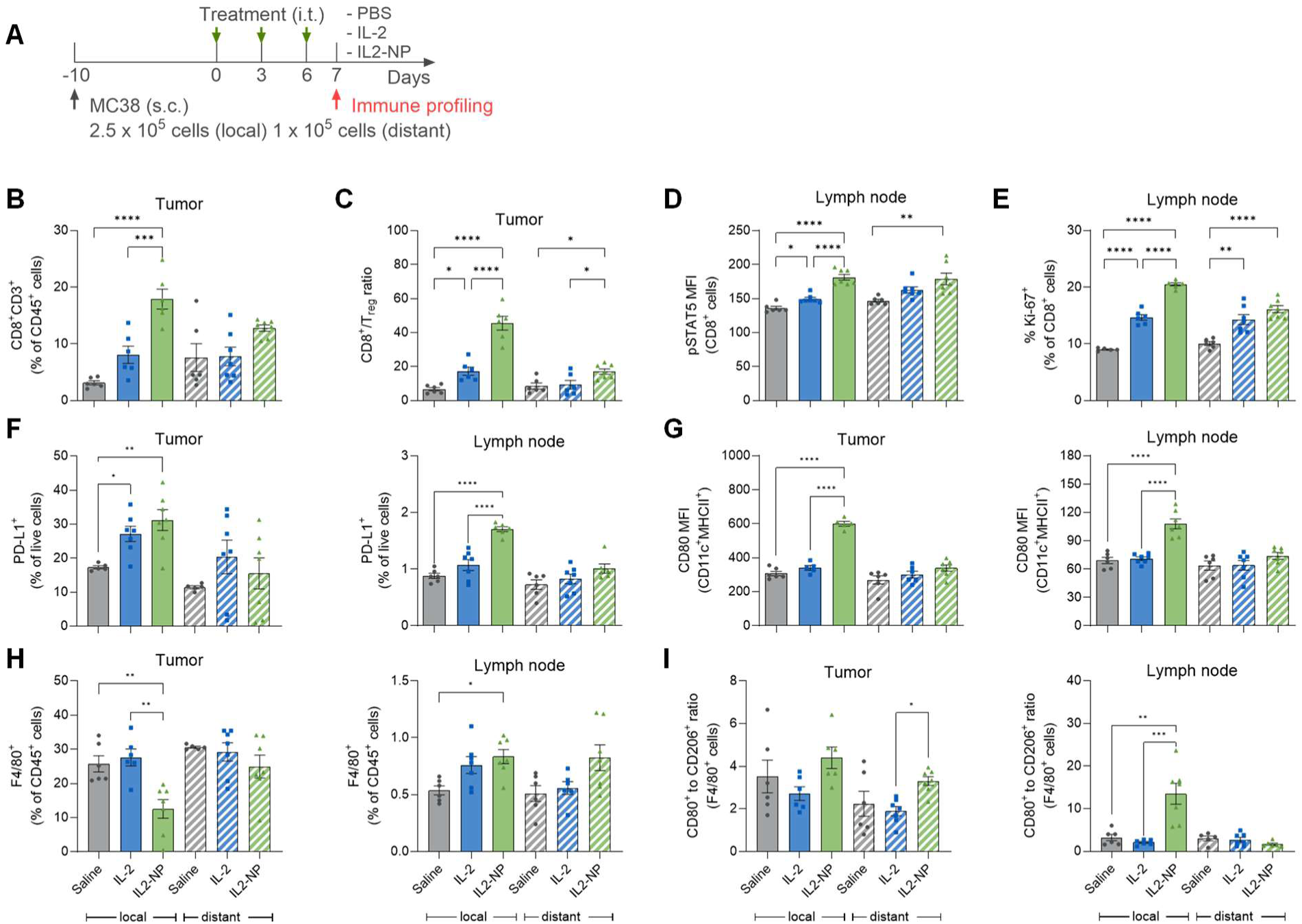
Immune remodeling of tumors and tumor-draining lymph nodes by IL2-NP versus free IL-2. (**A**) Schematics of in vivo immune profiling experiment in bilateral MC38 tumor model by flow cytometry. (**B**) Populations of CD8⁺ T cells in tumors. (**C**) Ratio CD8⁺ T cells and T_reg_ in tumors. (**D**) Phosphorylated STAT5 status of CD8^+^ T cells in tumor-draining lymph nodes. (**E**) Proliferation (Ki67^+^) status of CD8⁺ T cells in tumor-draining lymph nodes. (**F**) Percentage of PD-L1^+^ cells among live cells in tumors (left) and tumor-draining lymph nodes (right). (**G**) Maturation level (CD80) of CD11c^+^ MHCII^+^ dendritic cells in tumors (left) and tumor-draining lymph nodes (right). (**H**) Macrophage population (F4/80^+^) in tumors (left) and tumor-draining lymph nodes (right). (**I**) Ratio of “M1-like” pro-inflammatory macrophages (CD80^+^F4/80^+^) over “M2-like” anti-inflammatory macrophages (CD206^+^F4/80^+^) in tumors (left) and tumor-draining lymph nodes (right).

IL-2-associated signaling was assessed by measuring phosphorylated STAT5 (pSTAT5, Tyr694), a downstream readout of IL-2R activation, in T-cell subsets. IL2-NP treatment significantly increased pSTAT5 levels in CD8^+^ T cells in the tumor draining lymph nodes compared to both free IL-2 and saline controls (**Figure 5D**), and elevated pSTAT5 in tumors compared to saline (**Supplementary Figure S1B**). In contrast, no significant change in pSTAT5 was observed in T_reg_ in either tumor or draining lymph nodes (**Supplementary Figure S1C**). We next examined T-cell proliferation by Ki67 staining. Consistent with enhanced IL-2 signaling, IL2-NP treatment significantly increased CD8^+^ T cell proliferation compared to free IL-2 and saline controls in tumor-draining lymph nodes (**Figure 5E**), while T_reg_ proliferation remained unchanged (**Supplementary Figure S1D**). Notably, these immune changes were more pronounced in local tumors than in distal tumors, whereas significant changes were observed in the draining lymph nodes of both tumors. This pattern may reflect the timing of analysis (1 day after last dose), with systemic immune changes detectable earlier in lymphoid tissues than in distant tumor tissues.

Additionally, IL2-NP treatment resulted in increased PD-L1 expression in live cells of local tumors and draining lymph nodes, with a pronounced increase compared to free IL-2 treatment in the local tumor-draining lymph nodes (**Figure 5F**). The increased PD-L1 expression may reflect an IFN-γ driven immune resistance response following T cell activation.^34^

The changes in myeloid cell populations associated with IL2-NP treatment were assessed. Although IL2-NP treatment did not significantly alter the dendritic cell abundance, it significantly increased activation (CD80 expression) of dendritic cells in tumors and tumor-draining lymph nodes (**Figure 5G**). IL2-NP treatment also decreased macrophage (F4/80^+^) population in treated tumors and increased this population in local draining lymph nodes (**Figure 5H**). Additionally, IL2-NP significantly increased the ratio of “M1-like” (CD80^+^F4/80^+^) to “M2-like” (CD206^+^F4/80^+^) macrophages in distant tumors and local tumor-draining lymph nodes (**Figure 5I**), suggesting a shift toward a more pro-inflammatory, antitumor macrophage phenotype.

Together, these findings show that IL2-NP treatment was associated with enhanced CD8^+^ T cell signaling and proliferation, an increased CD8/T_reg_ ratio, increased PD-L1 expression and favorable changes in myeloid cell phenotype in the tumors and tumor-draining lymph nodes.

### 2.5. IL2-NP treatment was efficacious in ovarian cancer and triple-negative breast cancer models

Next, we evaluated whether the therapeutic activity of IL2-NP extended beyond the MC38 colorectal cancer model to tumor models with distinct immune barriers. The ID8 ovarian cancer and E0771 breast cancer models were selected because ID8 is characterized by low T-cell infiltration and high levels of immunosuppressive myeloid cells,^35,36^ and E0771 has been reported to exhibit adaptive resistance to PD-1 blockade.^37–39^

We first evaluated IL2-NP in the ID8-luc peritoneal metastasis model of ovarian cancer. In this model, luciferase-expressing ID8 ovarian cancer cells were injected intraperitoneally, resulting in tumor nodule formation in the peritoneal cavity and eventual development of ascites, recapitulating clinical progression of advanced ovarian cancer.^36,40^ Tumor burden was monitored by in vivo bioluminescence imaging (IVIS) and the experimental timeline is shown in **Figure 6A**. Combination treatment with ICIs and IL2-NP reduced the tumor burden (**Figure 6B**), prolonged overall survival (**Figure 6C**) and delayed the onset of ascites (**Figure 6D**) compared with ICIs alone. Notably, this combination outperformed the combination of free IL-2 and ICIs, which did not improve the therapeutic outcome compared to ICIs alone.

**Figure 6:**
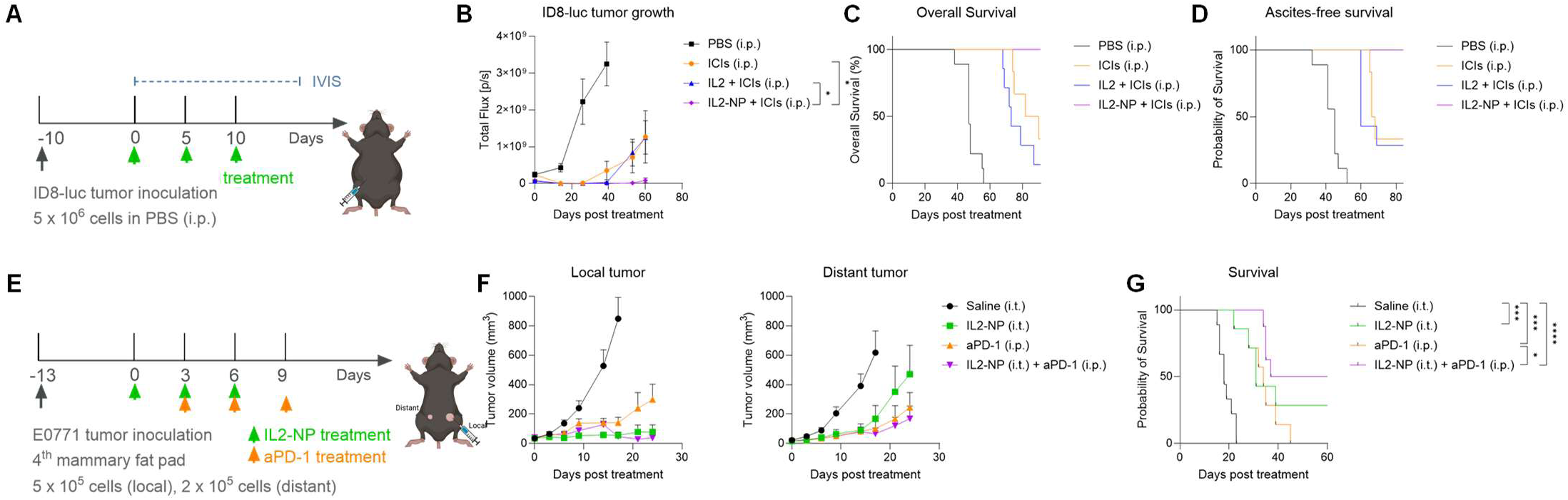
Efficacy of IL2-NP in a metastatic ID8 ovarian cancer model and a bilateral orthotopic E0771 breast tumor model. (**A**) Study timeline of the ID8-luc peritoneal metastasis model. (**B**) ID8-luc tumor growth determined by average photon flux via bioluminescence imaging using IVIS. (**C**) Kaplan–Meier Survival curve. (**D**) Ascites-free survival curve (duration without development of ascites following tumor inoculation). (**E**) Study timeline of the bilateral E0771 tumor model. (**F**) Tumor growth of the local treated E0771 tumor (left) and distant tumor (right). (**G**) Kaplan–Meier Survival curve.

We next evaluated IL2-NP in the E0771 model of triple-negative breast cancer, which has been reported to respond to anti-PD-1 initially but subsequently develop resistance.^37–39^ In the bilateral orthotopic E0771 model, the local tumor was treated by intratumoral injections of IL2-NP, and the contralateral distant tumor was left untreated to assess systemic antitumor response (**Figure 6E**). Local IL2-NP treatment reduced the growth of both local and distal tumors **Figure 6F**) and prolonged survival of the mice compared to saline control (**Figure 6G**). While this model showed moderate responsiveness to anti-PD-1 monotherapy, resistance ultimately developed and all mice reached the euthanasia endpoint due to tumor burden (**Figure 6G**). However, combining anti-PD-1 with IL2-NP significantly improved therapeutic outcomes and prolonged survival (**Figure 6F-G**). Together, these findings demonstrate that IL2-NP exhibits broad therapeutic efficacy, particularly when combined with immune checkpoint blockade.

## 3. Discussion

We report a new strategy for delivering wild-type IL-2 at high surface density using our patented nanoparticle platform. The data showed that increasing IL-2 surface density to approximately 8,000 molecules per particle promoted high-avidity engagement of IL-2 receptors, resulting in preferential expansion of CD8⁺ T cells without a proportional increase in T_reg_ expansion. This formulation enhanced IL-2 bioactivity, protected IL-2 from protease degradation, enhanced cellular uptake to T cells, and prolonged tumor retention of IL-2. In a mouse model of colorectal cancer, the IL2-NP treatment improved the control of both locally injected tumors and untreated contralateral tumors, and demonstrated a systemic anti-tumor immune response that was dependent on CD8^+^ T cells. When IL2-NP was combined with immune checkpoint inhibitors, the treatment led to complete tumor regression and development of tumor-specific immune memory that prevented tumor rechallenge. Together, these findings support our premise that the surface density of IL-2 on our nanoparticles can be tuned to achieve desirable outcomes.

While many IL-2 therapeutics currently in development focus on extending half-life and/or reducing T_reg_ activation by blocking IL-2Rα engagement, these approaches may alter native IL-2 biology that is important for eliciting effective antitumor immunity. For instance, recent evidence suggests that tumor-reactive CD8⁺ T cells express IL-2Rα and benefit from intact IL-2Rα signaling, raising the possibility that some engineered IL-2 variants may compromise therapeutically relevant IL-2 biology.^20,21^ In contrast, our strategy preserves native wild-type IL-2 while altering only how it is presented to relevant cells within the tumor. Specifically, at the same total IL-2 concentration, nanoparticles with higher IL-2 density preferentially enhanced CD8^+^ T cell expansion relative to lower-density counterparts. This observation supports the concept that multivalent presentation (avidity) of IL-2 can increase functional engagement of IL-2 receptors on CD8^+^ T cells. Prior work has shown that IL-2 receptor clustering at the cell surface influences receptor endocytosis and downstream signaling.^41^ One study demonstrated that increasing the surface density of IL-2 on hydroxyethyl starch nanocapsules enhanced T-cell uptake, whereas lower IL-2 density preferentially favored T_reg_ uptake.^42^ Still, the reported maximum surface density achieved (1,660 IL-2 molecules per particle) and/or the conjugation strategy may not have been optimal, as no enhancement in activity was observed in the IL-2-dependent CTLL-2 cell line relative to free IL-2.

Further, nanoparticle-based strategies have been explored to improve the therapeutic index of IL-2 by enhancing tumor retention and limiting systemic exposure. For example, BALLkine-2 utilizes micron-sized mesoporous silica particles to load IL-2 within their pores, enabling sustained local release following peritumoral injection.^43^ Other approaches have used liposomes with surface conjugated IL-2,^44,45^ or lipid nanoparticles encapsulating IL-2 mRNA.^46^ These studies also highlight design tradeoffs such as nanoparticle size, surface charge, cargo loading density, and cytokine presentation format. For example, relatively large nanoparticle size or strong positive surface charge can promote non-specific cell uptake and phagocytic clearance, while a net negative surface charge may hinder receptor engagement.^47^ Additionally, conjugation strategy may alter the IL-2 activity compared to electrostatic loading of IL-2 and often yield lower loading density. Lastly, releasing of IL-2 prior to receptor engagement does not take advantage of binding avidity. Our IL2-NP platform is distinguished by the high-density presentation of IL-2 on the nanoparticle surface and its ability to protect IL-2 from enzymatic degradation by the NP’s PEG layer without significantly interfering with IL-2 receptor engagement. The net charge is near neutral and the IL-2 electrostatic loading is strong enough that it does not release IL-2 prior to receptor engagement. Lastly, the prolonged tumor retention of this hybrid polymeric–inorganic nanoparticle represents an important advantage over lipid nanoparticles.

The in vivo immune profiling results provide further insights into how high-density wild-type IL-2 presentation by nanoparticle promotes immune changes that favor antitumor response. Compared to free IL-2, IL2-NP showed increased binding to CD8^+^ T cells, increasing their STAT5 phosphorylation and proliferation. IL2-NP increased the abundance of CD8^+^ T cells without increasing T_reg_, resulting in a higher CD8^+^/T_reg_ ratio in both local and distant tumors. Beyond its effect on T cells, IL2-NP treatment was associated with increased dendritic cell maturation and increased ratio of M1-like over M2-like macrophages in tumor and tumor-draining lymph nodes. Together, these findings suggest that IL2-NP may act not only through direct effects on T cells, but also through broader remodeling of the tumor microenvironment from an immunosuppressive state (**Figure 7A**) toward a more immunostimulatory state (**Figure 7B**).

**Figure 7:**
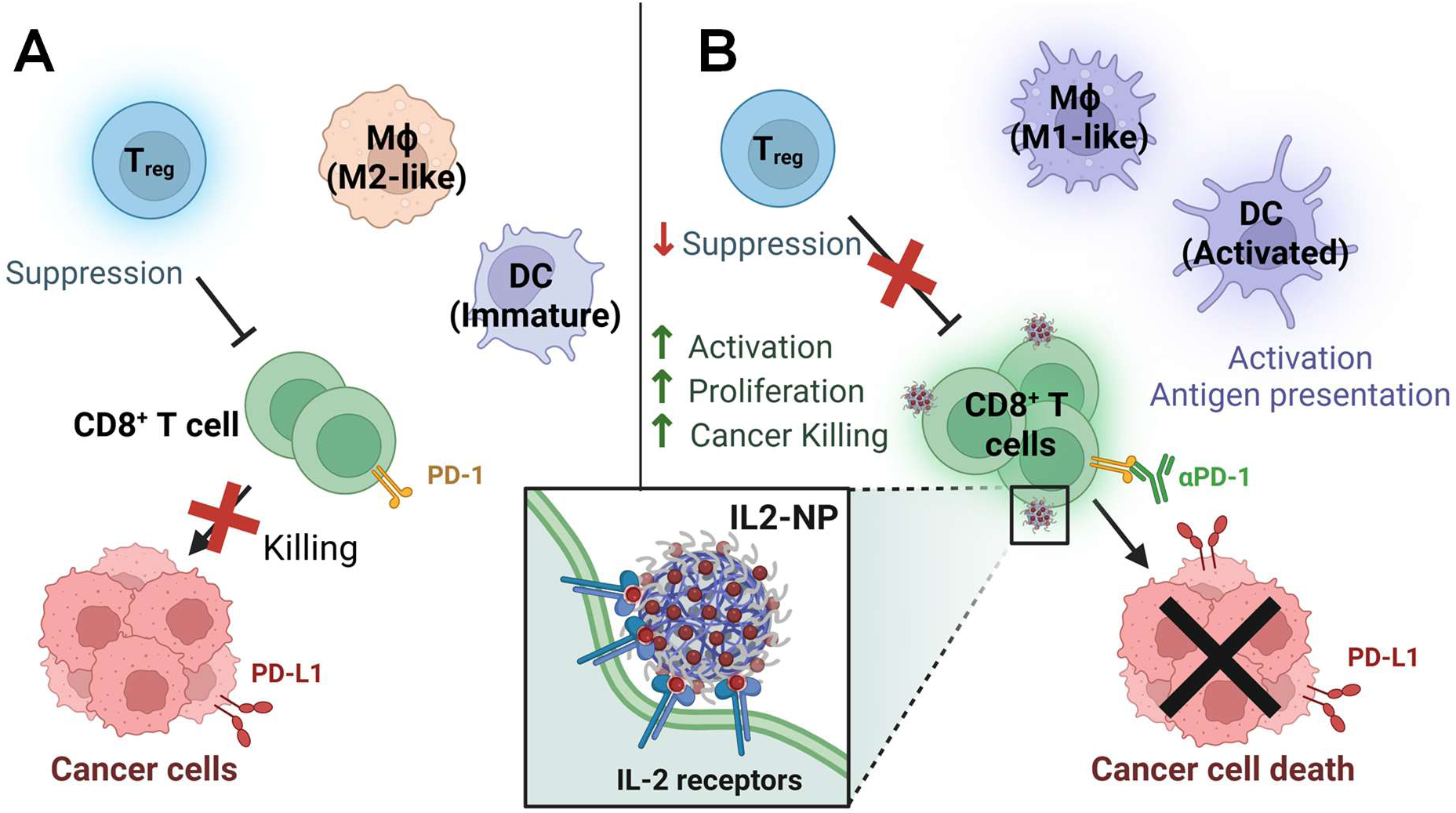
Proposed mechanism of action of IL2-NP. (**A**) In the untreated tumor microenvironment, regulatory T cells (T_reg_), immature dendritic cell (DC), “M2-like” anti-inflammatory macrophages (MΦ) suppress cytotoxic (CD8^+^) T cell activity and limit anti-tumor immunity. (**B**) IL2-NP treatment enhances CD8^+^ T cell activation and expansion through high-avidity binding to IL-2 receptors (inset) without increasing T_reg_ population, increases DC maturation and macrophage polarization toward a more pro-inflammatory “M1-like” phenotype. Overall, these changes shift the tumor immune microenvironment from immunosuppressive to immunostimulatory to enhance anti-tumor immunity and synergize with immune checkpoint inhibitors. (Created with BioRender.com)

These findings are meaningful because dendritic cells and macrophages are critical antigen-presenting cells that can support tumor-specific CD8^+^ T cell priming,^48^ whereas tolerogenic dendritic cells and immunosuppressive tumor-associated macrophages are known to support tumor growth, metastasis, and T_reg_.^49^ Still, the findings should be interpreted as treatment-associated changes rather than evidence for a defined causal pathway. For example, dendritic cell recruitment and maturation have been attributed to cytokines released by lymphocytes upon stimulation.^14,30^

Another important finding was the enhanced antitumor activity of IL2-NP when combined with immune checkpoint inhibitors. In the MC38 colorectal tumor model, combining IL2-NP with ICIs produced complete responses in all treated mice, whereas ICIs alone were less effective. Improved efficacy with IL2-NP + ICIs was also observed in the ID8-luc ovarian peritoneal metastasis model, where ICIs alone or ICIs combined with free IL-2 were less effective. Similarly, in the E0771 breast tumor model, IL2-NP significantly improved the antitumor efficacy of anti-PD-1 therapy. These findings support the premise that IL2-NP may help address key limitations of immune checkpoint blockade, such as insufficient quantity or functional activity of T cells within the immunosuppressive tumor microenvironment. The observed PD-L1 upregulation after IL2-NP treatment indicates treatment-induced immune activation and may reflect a state in which tumors are more susceptible to PD-1/PD-L1 blockade treatment.^34^ In addition to augmenting immune checkpoint blockade, we believe the IL2-NP will significantly befits the rapidly growing fields of cancer vaccines and tumor-infiltrating lymphocyte (TIL) therapies, as both require effective and safe strategies for expanding CD8⁺ T cells.

Lastly, this study also highlights the potential value of locoregional cytokine delivery. Because recombinant IL-2 has a short circulation half-life, systemic IL-2 therapy typically requires repeated high-dose administration (600,000 IU/kg IV every 8 hours, maximum 14 doses)^12^ to achieve robust antitumor immunity, but these regimens are frequently limited by dose-dependent toxicities.^15^ The same requirement is seen preclinically; achieving antitumor activity with free WT IL-2 in mice require intensive repeated systemic dosing, such as 10 µg given i.p. twice daily for 20 doses (∼200 µg cumulative),^43^ or 3 mg/kg once daily for five days (∼300 µg cumulative),^50^ to sustain high IL-2 exposure. By contrast, intratumoral IL2-NP administered every 3 days for three doses (30 µg total) increases local tumor IL-2 exposure, drives the expansion of tumor-specific T cells that traffic to distant tumor sites, and minimizes systemic IL-2 exposure, highlighting the potency of this formulation and the feasibility of its intratumoral administration.

## 4. Conclusion

In summary, high-density surface presentation of wild-type IL-2 on polymer-coated mesoporous silica nanoparticles preferentially expanded CD8^+^ T cells over T_reg_, protected IL-2 from proteolysis, and prolonged intratumoral retention. Local IL2-NP treatment remodeled tumor microenvironment and drove systemic CD8^+^ T cell dependent antitumor immunity, producing durable responses with immunological memory when combined with immune checkpoint blockade, with efficacy extending across colorectal, ovarian and triple-negative breast cancer models. These findings suggest that the mode of cytokine presentation is a key yet underexplored aspect of cytokine delivery. Beyond IL-2, the modular platform and tunable surface presentation may be broadly applicable to other cytokines and biologics.

## 5. Experimental

### 5.1. Nanoparticle synthesis and characterization

Polymer coated MSNPs were synthesized and characterized similarly to published protocols.^22^ The MSNPs have a uniform size of about 50 nm in diameter as determined by transmission electron microscopy (TEM). Polymer composition and the mass concentration of NP was determined by thermogravimetric analysis (TGA Q50, TA Instruments) by heating 50 µL of sample to 650°C (20°C/min). Particle concentration was measured by nanoparticle tracking analysis using Zetaview instrument (Particle Metrix, Germany). For IL-2 loading, mouse recombinant IL-2 (PeproTech/Thermo Fisher, cat. #212-12) was reconstituted according to the manufacturer’s recommendation and added to NP at 0.3-2.0 wt.% of NP, and mixed on an orbital shaker at 350 rpm overnight at 4°C. Final concentration of NP was 10 mg/mL in PBS pH 7.2 (Gibco, cat. #20012050), and the IL-2 loaded NP (IL2-NP) was stored at -80°C until further use. IL-2 loading was determined by quantification of unbound IL-2 in the supernatant using Pierce BCA protein assay (ThermoFisher, cat. #A65453). Hydrodynamic size and zeta potential of nanoparticles were measured using Zetasizer (Malvern, ZS-90). Nanoparticle at 10 mg/mL in PBS pH 7.2 was diluted to 0.2 mg/mL in PBS for size measurement, and diluted 200-fold in DI water for zeta potential measurement, respectively. IL-2 was tagged with Alexa Fluor 647 NHS Ester (Invitrogen, cat. #A37573) or Alexa Fluor 680 NHS Ester (Invitrogen, cat. #A37574) at 1:1 molar ratio. The mixture was shaken at room temperature for 2 hours and the excess dye was removed using 3kDa MWCO Amicon ultra centrifugal filter (MilliporeSigma, cat. #UFC500308). Dye-tagged IL-2 was loaded on NP as described previously.

### 5.2. Drug release profile assessment by ELISA

IL2-NP or free IL-2 was diluted with PBS pH 7.4, or 100 mM sodium acetate buffer pH 4.5 or pH 5.5 containing 0.1% bovine serum albumin to the final concentration of 1 mg/mL NP and 0.02 mg/mL IL-2. The mixture was shaken on an orbital shaker at 100 rpm and 37°C. On days 0, 4, and 7 days post incubation, samples were centrifuged to remove the nanoparticles. Released IL-2 in the supernatant was quantified using mouse IL-2 ELISA (BioLegend, cat. #431004) following the manufacturer’s protocol. IL-2 release was calculated as the ratio of IL-2 in the supernatant to free IL-2 (without nanoparticles).

### 5.3. Cell culture method and primary cell isolation

CTLL-2 and E0771 cell lines were purchased from ATCC. CTLL-2 was maintained in RPMI-1640 (Gibco, cat. #11875119) with 2 mM L-glutamine, 1 mM sodium pyruvate, 10% FBS (Gibco, cat. #A5669801) and 10% T-STIM with Con A (Corning, cat. #354115). E0771 was maintained in DMEM (Gibco, cat. #11965118) with 10% FBS and 10 mM HEPES (Gibco, cat. #15630080). MC38 cell line was purchased from Kerafast (cat. #ENH204-FP) and maintained in DMEM with 10% FBS, 2mM L-glutamine (Gibco, cat. #25030081), 0.1 mM nonessential amino acids (Gibco, cat. #11140050), 1 mM sodium pyruvate (Gibco, cat. #11360070), 10 mM HEPES, and 50µg/mL gentamycin sulfate (Gibco, cat. #15710064). ID8-luc cell line was purchased from Ubigene (cat. # YC-C103-Luc-P) and maintained in DMEM with 10% FBS. LLC-JSP cell line was gifted from Don Gibbons lab at MD Anderson Cancer Center and maintained in RPMI + 10% FBS.

Splenocyte harvest: spleen from six- to eight-week-old female C57BL/6 mice was cut into small sections and incubated in digestion media containing 1 mg/mL Collagenase D (Gibco, cat. #17104019) and 0.1 mg/mL DNase I (Sigma-Aldrich, cat. #10104159001) in HBSS (ThermoFisher, cat. #PI88284) at 37°C for 30 min and mechanically dissociated by passing through 70 µm cell strainers. Red blood cells were lysed using RBC lysis buffer following the manufacturer’s protocol (Invitrogen, cat. #50-112-9743), and the resulting immune cells were cultured in RPMI-1640 with 10% FBS and 1X P/S (Penicillin/Streptomycin, Gibco, cat. #15140122).

T cell isolation: T cells were isolated from dissociated splenocytes using EasySep Mouse T Cell Isolation Kit (STEMCELL Technologies, cat. #19851) following the manufacturer’s protocol, and cultured in RPMI-1640 with 10% FBS and 1X P/S or cryopreserved in FBS with 7.5% DMSO.

### 5.4. Cell culture assays

T cells were seeded at 5 × 10^5^ cells/mL in RPMI + 10% FBS (without IL-2) for 2-4 hours and treated with IL2-NP or free IL-2 for 48 hours. Cell viability was determined by CellTiter-Glo assay (Promega, cat. #G9242) following the manufacturer’s protocol.

Co-culture assay: isolated splenic T cells from six- to eight-week-old female C57BL/6 mice were pre-activated with plate coated anti-CD3 antibodies (1 µg/mL) and soluble anti-CD28 (5 µg/mL) antibodies for 48 hours. Then, 3 x 10^4^ activated T cells were co-cultured with 3 x 10^3^ LLC-JSP mouse lung cancer cells in 96-well plates and treated with PBS, free IL-2, IL2-NP (0.2 µg/mL IL-2 concentration) or NP without IL-2. Cell culture media was collected 24 hours later for analysis of IFN-γ and Granzyme B by ELISA following the manufacturer’s protocols (ThermoFisher, cat. #88-7314 and #88-8022).

Confocal imaging: one million CTLL-2 cells were suspended in 1 mL RPMI media with 10% FBS, and treated with nanoparticles loaded with AF647 dye-tagged IL-2 (AF647-IL2-NP, 50 µg/mL NP, 0.6 µg/mL IL-2) or soluble AF647-IL2 (3 µg/mL IL-2) for one hour at 37°C with rotary mixing. Cells were washed with PBS, stained with CellBrite 488 (Biotium, cat. #30090) and Hoechst 33342 (Invitrogen, cat. #H3570) following the manufacturer’s protocol, and fixed with 4% paraformaldehyde at room temperature for 20 minutes. The fixed cells were washed with PBS and attached to 8-well chambered coverslip (ibidi, cat. #80806) at 300 µL/well at 4°C overnight, and the coverslip was imaged on Zeiss LSM 880 confocal microscope.

### 5.5. Syngeneic mouse tumor models and treatments

Animals were recruited and used under approved IACUC protocol of OHSU. Six- to eight-week-old female NCI C57BL/6NCr mice from Charles River Laboratories were inoculated with MC38 cells on the flank (2.5 × 10^5^ cells on local and 1.0 × 10^5^ cells on distant sites), E0771 cells on 4^th^ mammary fat pads (5.0 × 10^5^ cells on local and 2.0 × 10^5^ cells on distant sites), or ID8-luc cells by intraperitoneal injection (5.0 × 10^6^ cells). Mice were randomized by tumor volume or tumor burden by IVIS before treatment.

In bilateral MC38 and E0771 tumor models, local tumors were treated every three days for three doses with IL2-NP (0.5 mg NP, 10 µg IL-2) or free IL-2 at equivalent dose by intratumoral injection 7 to 13 days post tumor inoculation. In indicated groups, mice were also treated with PD-1 Ab (200 µg per mouse, clone RMP1-14, BioXCell, cat. # BE0146) and/or CTLA-4 Ab (100 µg per mouse, clone 9H10, BioXCell, cat. #BE0131) by i.p. injection every three days for three doses. Tumors were measured with Vernier Caliper and volume calculated by V = 0.5 x length x width^2^. In CD8 depletion study, mice were i.p. injected with CD8 depleting antibody (200 µg per mouse, clone 53-6.7, BioXCell, cat. #BE0004-1) one day prior to initiation of treatment, twice a week for two weeks.

In metastatic ID8-luc tumor model, mice were treated with IL2-NP (0.5 mg NP, 10 µg IL-2) by i.p. injection on days 10, 15, and 20 post tumor inoculation, and PD-1 antibody on days 15, 20, and 25. For tumor burden monitoring, mice were injected i.p. with 150 mg/kg D-Luciferin, Potassium Salt (Gold Biotechnology, cat. #LUCK-1G) in 200 µL PBS 20 minutes prior to IVIS imaging under isoflurane anesthesia.

### 5.6. Flow cytometry and immune profiling

In immune profiling study, mice were euthanized 7 days post initiation of treatment (one day after the last dose), tumors and tumor-draining lymph nodes were harvested and processed for flow cytometry analysis similar to published protocols.^23,51^ Briefly, tumors were cut into small pieces, incubated in digestion media at 37°C for 30 min and mechanically dissociated by passing through 70 µm cell strainers. Red blood cells were lysed using RBC lysis buffer (Invitrogen, cat. # 00-4300-54). Cells were stained with Live/Dead fixable blue (Invitrogen, cat. #L34962) following the manufacturer’s protocol. FcR was blocked with anti-mouse Fc Shield (2.4G2, Tonbo Biosciences, cat. #70-0161) at 4°C for 15 min. Surface molecules were stained with fluorophore conjugated antibodies diluted in Brilliant Stain Buffer (BD, cat. #563794) for 30 min at 4°C. Cells were washed with FACS buffer containing 1% BSA (Fisher Scientific, BP9703100) in PBS pH 7.2, and fixed with 4% paraformaldehyde in PBS for 20 min at 4°C. Intracellular markers were stained using BD Cytofix/Cytoperm Fixation/Permeabilization Kit (BD, cat. #554714), or Transcription Factor Phospho Buffer Set (BD, cat. #563239) following the manufacturer’s protocol. Stained cells were run on BD FACSymphony A5 Cell Analyzer (BD Biosciences) or Cytek Aurora spectral cytometer (Cytek Biosciences) and analyzed using FlowJo. In ex vivo T cell experiments, isolated T cells were rested in RPMI-1640 with 10% FBS and 1X P/S for 2-4 hours. Cells were then treated and harvested at the time indicated in figure legends. The cells were washed with PBS and stained with Live/Dead fixable blue (Invitrogen, cat. #L34962) following the manufacturer’s protocol. Staining and analysis followed immune profiling study above.

The following antibodies were purchased from BioLegend: BV421 anti–mouse MHC II (I-A/I-E) (M5/114.15.2, cat. #107632); BV605 anti–mouse CD80 (16-10A1, cat. #104729); BV711 anti–mouse CD8 (53-6.7, cat. #100759); BV785 anti–mouse NK1.1 (PK136, cat. #108749); BV785 anti–mouse CD4 (RM4-5, cat. #100552); FITC anti–mouse CD3 (17A2, cat. #100204); FITC anti–mouse CD19 (6D5, cat. #115506); PerCP-Cy5.5 anti– mouse CD11c (N418, cat. #117328); PE-Cy7 anti–mouse PD-1 (29F.1A12, cat. #135216); PE-Cy7 anti–mouse CD206 (C068C2, cat. #141720); AF647 anti–mouse FoxP3 (MF-14, cat. #126408); AF647 anti–mouse F4/80 (BM8, cat. #123122); APC-Cy7 anti–mouse CD45 (30-F11, cat. #103116). The following antibodies were purchased from BD Biosciences: PerCP-Cy5.5 anti–mouse CD4 (RM4-5, cat. #550954); PE anti–mouse phospho-STAT5 (47/Stat5[pY694], cat. #612567); PE anti–mouse PD-L1 (MIH5, cat. #558091);.efluor450 anti– mouse Ki-67 (GB11, cat. #48-5698-82) was purchased from Invitrogen.

### 5.7. Tumor retention analysis

Mice bearing bilateral MC38 tumors were treated with intratumoral injections of saline, AF680-IL2 or AF680-IL2-NP to the local tumors. One day later, the tumors were harvested, homogenized and the AF680 fluorescence intensity was measured using LI-COR Odyssey imaging system. AF680-IL2 concentration was quantified against the calibration curve of AF680-IL2 or AF680-IL2-NP diluted in tumor tissue homogenate. Mass of AF680-IL2 in tumor was calculated based on the AF680-IL2 concentration and tumor volume, and percentage of injected dose was calculated based on injected dosage.

### 5.8. Serum clinical chemistry

Blood was collected via cardiac puncture after euthanasia and the whole blood was rested at room temperature to clot for 30 min to 1 hour. Serum was isolated by removing the clot by centrifugation at 1,500 x g for 10 minutes at 4°C, and stored at -20°C until analysis. The serum biochemistry parameters were analyzed on Beckman AU680 by IDEXX BioAnalytics (CA, USA).

### 5.9. Statistical analysis and AI use

GraphPad Prism was used for statistical analysis. In vitro data are presented as mean ± SD, whereas in vivo data are presented as mean ± SEM. Comparison between two groups was performed using t-test. Comparison between more than two groups was performed with one-way ANOVA with Tukey’s correction for multiple comparisons. Comparison in longitudinal tumor studies was performed with mixed model Two-Way repeated measures ANOVA with Tukey’s correction for multiple comparisons. Kaplan–Meier survival curve was analyzed with Log-rank Mantel–Cox test. ns: not significant, *P ≤ 0.05, **P ≤ 0.01, ***P ≤ 0.001, ****P ≤ 0.0001. Artificial intelligence-generated content (AIGC) tools (e.g., ChatGPT) were used solely for English language editing and grammatical improvement.

### 5.10. Use of artificial intelligence

Artificial intelligence-generated content (AIGC) tools (e.g., ChatGPT) were used for English language editing and grammatical improvement.

## 6. Supplementary figure

**Supplementary Fig. S1:**
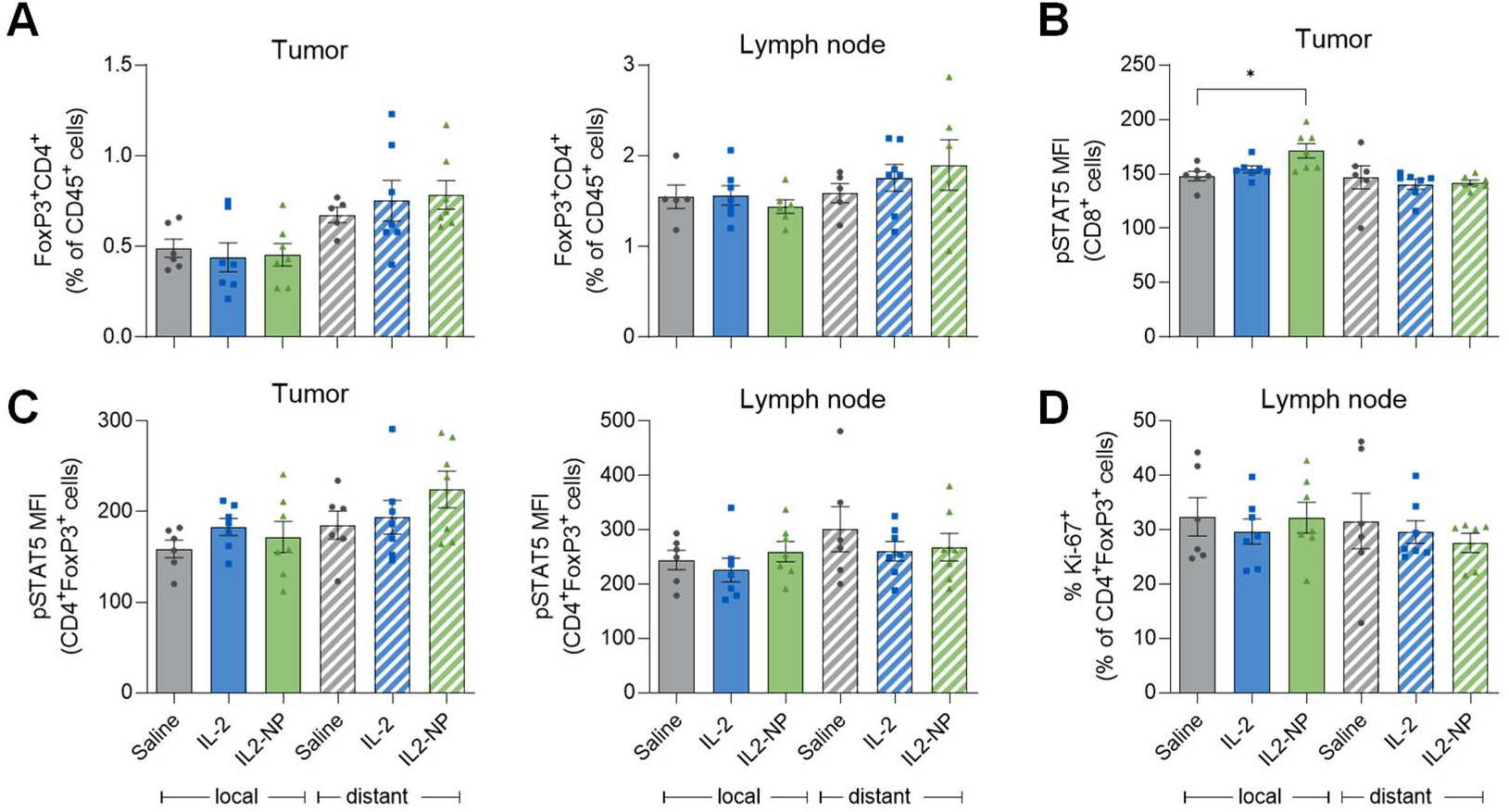
Analysis of CD8^+^ T cells and T_reg_ in mice from Fig. 4E. (**A**) Population of T_reg_ (CD4^+^FoxP3^+^) in tumors (left) and tumor-draining lymph nodes (right). (**B**) Phosphorylated STAT5 status of CD8^+^ T cells in tumors. (**C**) Phosphorylated STAT5 status of T_reg_ (CD4^+^FoxP3^+^) in tumor-draining lymph nodes. (**D**) Proliferation (Ki67^+^) status of T_reg_ in tumor-draining lymph nodes.

## Acknowledgement

This work was funded by the NCI R44CA265751, R44CA265752, R44CA285233, Women’s Health Circle of Giving at OHSU, Frohnmayer Hicks Sciarretta Cancer Research Scholar award from the Knight Cancer Institute, Joe W. and Jane E. Gray Professorship Fund, and Chulabhorn Royal Academy (by National Science Research and Innovation Fund, FRB660044/0240, Project code 180868). Its contents are solely the responsibility of the authors and do not necessarily represent the official views of the NIH and US government. We thank Natalie White for assistance in core nanoparticle synthesis, and Lindsey Savoy for assistance with the ELISA assay setup and data collection. We thank Dr. Xiaolin Nan and Dr. Megan Burger of OHSU’s Knight Cancer Institute for their valuable input and Dr. Xiaolin Nan for his independent reviewing of the data in this manuscript as required by OHSU’s conflict of interest guidelines. We thank Dr. Don Gibbons lab of MD Anderson Cancer Center for generous gift of LLC-JSP lung cancer cell line. We thank OHSU shared resources cores: Flow Cytometry Shared Resource (RRID: SCR_009974) and Advanced Light Microscopy Core (RRID: SCR_009961) for access to instrumentation and technical support.

The data that support the findings of this study are available from the corresponding author upon reasonable request.

